## Supplemental materials for "DeepETPicker: Fast and accurate 3D particle picking for cryo-electron tomography using weakly supervised deep learning"

### A. Supplementary Methods

**A.1 Preprocessing of Tomograms.** A zero-mean normalization of voxel values is carried out for each input tomogram as follows:

$$x^* = \frac{x - \mu}{\sigma} \tag{1}$$

where $x \in \mathbb{R}^{N \times N \times N}$ denotes the input tomogram, $\mu$ and $\sigma$ denote the mean value and standard deviation of $x$, respectively, and $x^*$ denotes the zero-mean normalized tomogram.

**A.2 Software Design.** DeepETPicker is open-source software implemented in Python with a user-friendly graphical interface (Supplementary Fig. 1). It integrates multiple functions, including picking of particles, visualization of annotated particles, pre-processing of input tomograms, generation of simplified masks, and configuration of parameters for training and inference. DeepETPicker picks 3D particles of varying sizes and structures from simulated and experimental cryo-ET datasets with the best overall speed and accuracy in comparing with competing state-of-the-art methods.

**A.3 Hyperparameter Setting**.

There are several hyperparameters ($t_g, t_{seg}, t_{dist}$, .etc) for DeepETPicker. Here, we discuss how the setting of these hyperparameters affects results of particle picking.

According to **Equations (1, 4)**, the hyperparameter $\boldsymbol{t_g}$ determines the shape of Gaussian masks ($mask_{gaussian}$). The mask of $mask_{gaussian}$ becomes a ball mask when $\boldsymbol{t_g} \geq \exp(-0.5) \approx 0.607$, and it becomes a cubic mask when $\boldsymbol{t_g} \leq \exp(-1.5) \approx 0.223$. Therefore, $mask_{gaussian}$ will be a Gaussian mask when $\boldsymbol{t_g} \in (0.223, 0.607)$. When $\boldsymbol{t_g}$ approaches 0.223, the generated Gaussian mask would be very similar to a cubic mask; When $\boldsymbol{t_g}$ approaches 0.607, the generated Gaussian mask would be very similar to a ball mask. To generate a Gaussian mask that differs sufficiently from ball/cubic masks, we choose the middle point between $-0.5$ and $-1.5$ and set $\boldsymbol{t_g} = \exp(-1) \approx 0.368$ in this study.

The output score maps of 3D-ResUNet are in the range of [0,1], in which the value of each voxel denotes its probability score of belonging to a certain class. $\mathbf{t_{seg}}$ is a selected threshold that transforms a soft score map into a binary map: a voxel with its value below $\mathbf{t_{seg}}$ is labeled as 0 and otherwise as 1 so that a binary map is generated. The influence of $\boldsymbol{t_{seg}}$ on the classification performance of DeepETPicker trained by different types of masks on SHREC2021 dataset is summarized in **Supplementary Table A1**. The results show that $\boldsymbol{t_{seg}}$ has little effect on the classification performance when it varies from 0.1 to 0.9. Therefore, we set the default value of $\boldsymbol{t_{seg}}$ to 0.5 in this study.

**Supplementary Table A1 | Influence of $t_{seg}$ on classification performance of DeepETPicker trained by different types of masks on the SHREC2021 dataset.**

| Label Type | $t_{seg}$ | | | | | | | | |
|---|---|---|---|---|---|---|---|---|---|
| | 0.1 | 0.2 | 0.3 | 0.4 | 0.5 | 0.6 | 0.7 | 0.8 | 0.9 |
| Real Mask | 0.817 | 0.816 | 0.816 | 0.815 | 0.815 | 0.815 | 0.815 | 0.815 | 0.815 |
| Gaussian, d=9 | 0.830 | 0.830 | 0.830 | 0.830 | 0.830 | 0.831 | 0.831 | 0.830 | 0.830 |
| Cubic, d=9 | 0.827 | 0.827 | 0.828 | 0.828 | 0.828 | 0.828 | 0.828 | 0.827 | 0.828 |
| Ball, d=9 | 0.829 | 0.828 | 0.828 | 0.829 | 0.829 | 0.829 | 0.829 | 0.829 | 0.828 |

The hyperparameter $\boldsymbol{t_{dist}}$ is the threshold for the minimal Euclidean distance between two particles. Normally, $\boldsymbol{t_{dist}}$ is set to be half of the diameter of the particle $\left\lceil \frac{d}{2} \right\rceil$, where $\lceil\cdot\rceil$ denotes the round-up operation. If the Euclidean distance between two particles is lower than $\boldsymbol{t_{dist}}$, the two particles are considered the same.

The hyperparameter $\boldsymbol{t_{lm}}$ is a threshold determining whether a local maximum is a particle. In our study we set $\boldsymbol{t_{lm}}$ as a constant of 0.1. The hyperparameter $\boldsymbol{N}$ is the size of a subtomogram. It needs to be a multiple of 8. It is recommended that this value be no less than 64, and the default value is 72. The hyperparameter $\boldsymbol{pad\_size}$ is the padding size for the overlap-tile strategy. Usually, it ranges from 6 to 12, and the default value is 12. The hyperparameter $\mathbf{max}\_\boldsymbol{epoch}$ is the total number of training epochs.

The default value 60 is usually sufficient. The hyperparameter $\boldsymbol{batch\_size}$ is the number of samples processed before the model is updated. It is determined by the GPU memory. Reducing this parameter may be helpful if an out-of-memory error is encountered.

In summary, while $\boldsymbol{t_{dist}}$ is set to be a half of the particle diameter and $\boldsymbol{batch\_size}$ is determined by the GPU memory, other hyperparameters can always use their default values. A more detailed description for the choice of hyperparameters can be found in https://github.com/cbmi-group/DeepETPicker.

### A.4 Ablation Study for 3D-ResUNet Architectural Customizations

Coordinated convolution incorporates the spatial context of the input images into the convolutional filters, while image pyramid inputs preserves features of input images at different resolution levels, which may effectively improve the performance of convolutional neural networks. For validation, an ablation study for coordinated convolution and image pyramid inputs was carried out (**Supplementary Table A2**). We can observe that coordinated convolution or image pyramid inputs improve the mean F1-score by ~1.5% individually. They mainly improve the classification performance of tiny particles. When coordinated convolution and image pyramid inputs are added simultaneously, the mean F1-score of all complexes improves by 4.2%, and the mean F1-score of tiny complexes improves by 8.1%.

**Supplementary Table A2 | An ablation study for coordinated convolution and image pyramid inputs on the SHREC2021 dataset.** CC denotes coordinate convolution, and IP denotes image pyramid inputs. Tomograms 0 to 2 are used for training of DeepETPicker, tomogram 8 is used for validation and tomogram 9 is used for testing. In the ablation study, pad_size is set as 0, and data augmentation is not used.

| Ablation Study | | Mean F1-score of classification | | | | |
|---|---|---|---|---|---|---|
| CC | IP | Tiny | Small | Medium | Large | Total |
| | | 0.227 | 0.488 | 0.830 | 0.922 | 0.612 |
| √ | | 0.234 | 0.500 | 0.871 | 0.915 | 0.627 |
| | √ | 0.253 | 0.485 | 0.868 | 0.928 | 0.627 |
| √ | √ | 0.308 | 0.525 | 0.873 | 0.930 | 0.654 |

An ablation study is also carried out for channels of 3D-ResUNet. We find that 3D-ResUNet with channels of [8, 16, 24, 36] achieves comparable performance to that of [24, 48, 72, 108]. The model size of 3D-ResUNet with channels of [8, 16, 24, 36] is only 3.4 MBytes, which validates its architectural efficiency.

**Supplementary Table A3 | An ablation study for channels of 3D-ResUNet on the SHREC2021 dataset.** Tomograms 0 to 7 are used for training of DeepETPicker, tomogram 8 is used for validation and tomogram 9 is used for testing.

| Channels [c1, c2, c3, c4] | Model size | Localization performance | | | | Classification Performance |
|---|---|---|---|---|---|---|
| | | AD | Recall | Precision | F1-score | Mean F1-score |
| [24, 48, 72, 108] | 28M | 1.18 | 0.933 | 0.961 | 0.947 | 0.835 |
| [8, 16, 24, 36] | 3.4M | 1.217 | 0.927 | 0.944 | 0.935 | 0.829 |
| DeepFinder | 11M | 2.22 | 0.867 | 0.869 | 0.868 | 0.784 |

### A.5 Ablation Study for Loss functions

A detailed study of different types of "weak labels" shows that simplified masks with constant diameters can be used to replace particles with different diameters, achieving performance comparable to that of real masks. Using simplified masks with constant diameters as the training labels eliminates the problem of class imbalance and simplifies our selection of loss functions. We performed an ablation study for loss functions, demonstrating that different losses achieve similar picking performances when using simplified masks with constant diameters (**Supplementary Table A4**).

**Supplementary Table A4.** An ablation study for loss functions on the SHREC2021 dataset. Gaussian masks with constant diameter of 9 is used. Tomograms 0 to 2 are used for training of DeepETPicker, tomogram 8 is used for validation and tomogram 9 is used for testing.

| Loss function | Localization performance | | | | Classification |
|---|---|---|---|---|---|
| | AD | Recall | Precision | F1-score | mean F1-score |
| Dice | 1.34 | 0.902 | 0.971 | 0.935 | 0.786 |
| MSE | 1.417 | 0.908 | 0.978 | 0.942 | 0.773 |
| Focal | 1.373 | 0.911 | 0.977 | 0.943 | 0.759 |
| IoU | 1.304 | 0.881 | 0.985 | 0.93 | 0.767 |

### A.6 Ablation Study for Data Augmentation

The data augmentation used in our study is composed of two types of transformations, namely mirror transformation and spatial transformation (including random cropping, elastic deformation, scaling and rotation). An ablation study was carried out for these two types of transformations (**Supplementary Table A5**). We find that on the SHREC2021 dataset, mirror transformation substantially improves classification F1-score by 8.3% and slightly improves the localization F1-score by 1.0%. Spatial transformation improves the localization F1-score by 2% and classification F1-score by 2%. This is because the spatial transformation such as random rotation/cropping effectively increase the diversity of the tomogram training data. When mirror transformation and spatial transformation are used jointly, the mean F1-score of classification is improved by 9.5% and the F1-score of localization is improved by 3.4%. The ablation study described above indicates that mirror transformation can effectively improve the classification performance.

**Supplementary Table A5. An ablation study for different transformations on the SHREC2021 dataset.** Tomograms 0 to 2 are used for training of DeepETPicker, tomogram 8 is used for validation and tomogram 9 is used for testing.

| Mirror Transformation | Spatial Transformation | Localization performance | | | | Classification |
|---|---|---|---|---|---|---|
| | | AD | Recall | Precision | F1-score | mean F1-score |
| | | 1.513 | 0.859 | 0.947 | 0.901 | 0.691 |
| √ | | 1.372 | 0.891 | 0.932 | 0.911 | 0.774 |
| | √ | 1.469 | 0.878 | 0.968 | 0.921 | 0.711 |
| √ | √ | 1.34 | 0.902 | 0.971 | 0.935 | 0.786 |

### A.7 Threshold settings for DeepFinder and TM

For DeepFinder, it would generate a file with five columns, i.e., class label, x, y, z and cluster size. For the EMPIAR-10045 and EMPIAR-10499 datasets, the diameter of ribosomes is about 23~24 voxels. During the training stage, we use spheres with a radius of 11 as labels. For a sphere with a diameter of 23, its volume can be calculated as $V = \frac{4\pi r^3}{3} = \frac{4\pi \times 11.5^3}{3}$. In the inference stage, particles with a cluster size in the range of [0.1V, 2V] are selected as the final result for EMPIAR-10045. Particles with a cluster size in the range of [0.2V, 2V] are selected as the final result for EMPIAR-10499. For

EMPIAR-10651, we use spheres with a radius of 11 as the labels, and its volume can be calculated as $V = \frac{4\pi r^3}{3} = \frac{4\pi \times 11^3}{3}$. In the inference stage, particles with a cluster size in the range of [0.1V, 2V] are selected as the final result for EMPIAR-10651. For EMPIAR-11125, we use spheres with a radius of 7 as the labels, and its volume can be calculated as $V = \frac{4\pi r^3}{3} = \frac{4\pi \times 7^3}{3}$. In the inference stage, particles with a cluster size in the range of [0.1V, 4V] are selected as the final result for EMPIAR-11125.

For template matching, we use mainly the template matching function “dynamo_match” of Dynamo. There are two parameters that may affect particle selection. The parameter 'cr' (cone range) defines orientations that will be looked for inside a cone. In our experiment, we use the most typical value of 360 (sampling the full sphere). The parameter 'cs '(cone sampling) determines the scanning density inside the sphere. In our experiment, we use the most typical values of 30 (sampling the full sphere). It will generate tbl-format table files, where the tenth column shows the cross-correction coefficient. For each tomogram, we obtain a plot of the cross-correlation values found on the local maxima of the cc volume with the order. The cross-correlation values of the peaks appeared in an ascending order. We check the quality of the peaks by auxiliary clicking on the curve to select one particle and then selecting certain visualization option. We click on a couple of particles in the area of kink in the cross-correlation to roughly estimate the cross-correlation threshold. The detailed thresholds for TM for different datasets are provided in the following **Supplementary Table A6**.

**Supplementary Table A6. Cross-correlation threshold of TM for different datasets.**

| EMPIAR-10045 | | EMPIAR-10499 | |
|---|---|---|---|
| IS002_291013_005 | 0.182 | TS_77 | 0.060 |
| IS002_291013_006 | 0.188 | TS_78 | 0.060 |
| IS002_291013_007 | 0.163 | TS_79 | 0.062 |
| IS002_291013_008 | 0.150 | TS_80 | 0.065 |
| IS002_291013_009 | 0.160 | TS_81 | 0.050 |
| IS002_291013_010 | 0.170 | TS_82 | 0.053 |
| IS002_291013_011 | 0.157 | TS_84 | 0.057 |
| | | TS_85 | 0.059 |
| | | TS_87 | 0.061 |
| | | TS_88 | 0.063 |
| EMPIAR-10651 | | EMPIAR-11125 | |

| | | | |
|---|---|---|---|
| k2dft20s_14apra0006 | 0.440 | CB_02 | 0.250 |
| k2dft20s_14apra0011 | 0.440 | CB_29 | 0.240 |
| k2dft20s_14apra0023 | 0.440 | CB_59 | 0.260 |

#### A.8 Split of Training/Validation/Test Sets

In practice, a scheme commonly followed by structural biologists for particle picking is to manually label a small number of particles, use these particles for training the deep learning model selected for particle picking, and, finally, use the trained model to pick particles from all tomograms.

For EMPIAR-10045, a total of 150 particles are manually labeled along different consecutive z slices. For crYOLO, DeepFinder, and DeepETPicker, the first 135 sorted particles are used for training, and the remaining 15 particles are used for validation.

For EMPIAR-10651, a total of 142 particles are manually labeled along different consecutive z slices. For crYOLO, DeepFinder and DeepETPicker, the first 128 sorted particles are used for training, and the remaining 14 particles are used for validation.

For EMPIAR-10499, a total of 117 particles are manually labeled along different consecutive z slices. For crYOLO and DeepETPicker, the first 106 particles are used for training and the remaining 11 particles are used for validation. For DeepFinder, because its initial training using 106 particles fails to converge, we increase the number of manually labelled particles to 703, with the first 650 particles used for training and the remaining 53 particles used for validation.

For EMPIAR-11125, a total of 571 particles are manually labeled along different consecutive z slices. For crYOLO, DeepFinder, and DeepETPicker, the first 514 particles are used for training, and the remaining 57 particles are used for validation.

For all these four datasets, the coordinates of the manually picked particles are sorted in the order of z, y, and x from the smallest to the largest before split of the training and validation sets. This sorting strategy reduces the risk of overlap between the training and the validation sets.

In the inference stage, the trained model of each method is used to pick particles from all tomograms. For EMPIAR-10045 and EMPIAR-10499, to avoid overestimation of performance metrics, particles that are used initially for training and detected later in the inference stage are removed from subsequent data processing. However, for EMPIAR-10651 and EMPIAR-11125, only 3 tilt-series can be found and processed. Because the total numbers of particles detected by all methods are rather low, we do not remove the training particles so that enough particles can be used for subtomogram averaging. To check the level of performance metric overestimation, we have compared the precision-recall curves for the tested particle picking methods with and without the training particles on the EMPIAR-10651 and EMPIAR-11125 datasets in Supplementary Figure A1. Overall, it can be observed that the level of overestimation is relatively low and the conclusions on comparing performance of the tested methods remain qualitatively unchanged.

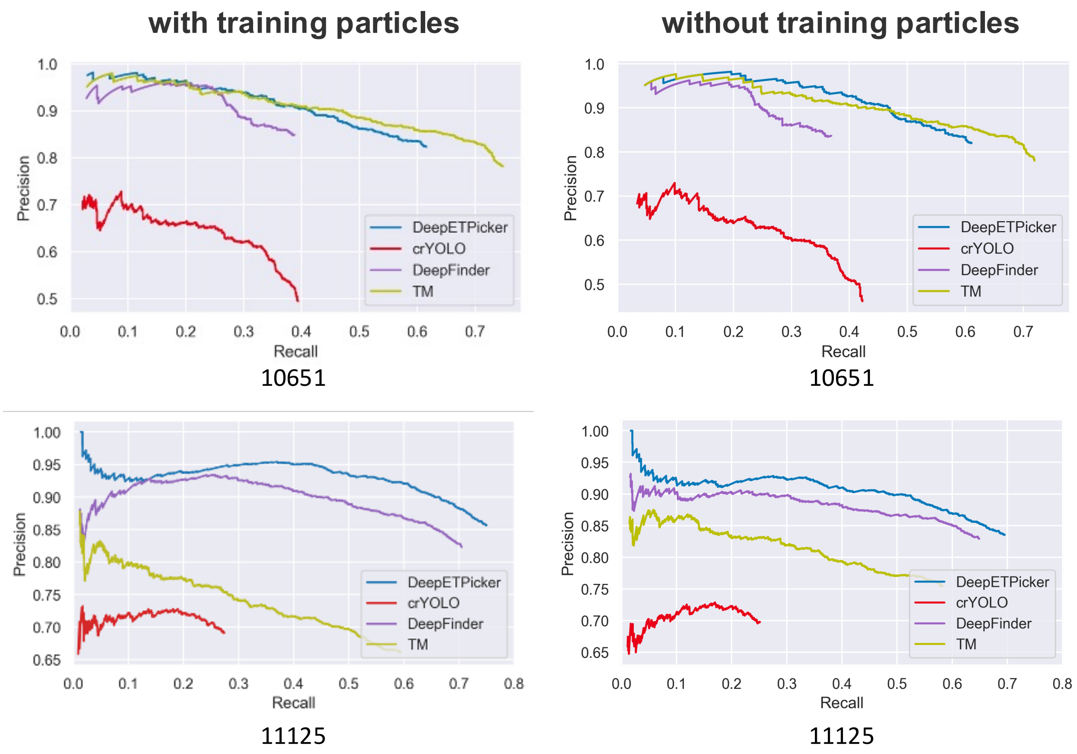


**Supplementary Figure A1. Precision-recall curves produced by different methods with and without training particles using manually picked particles as references.**

### B. Supplementary Figures

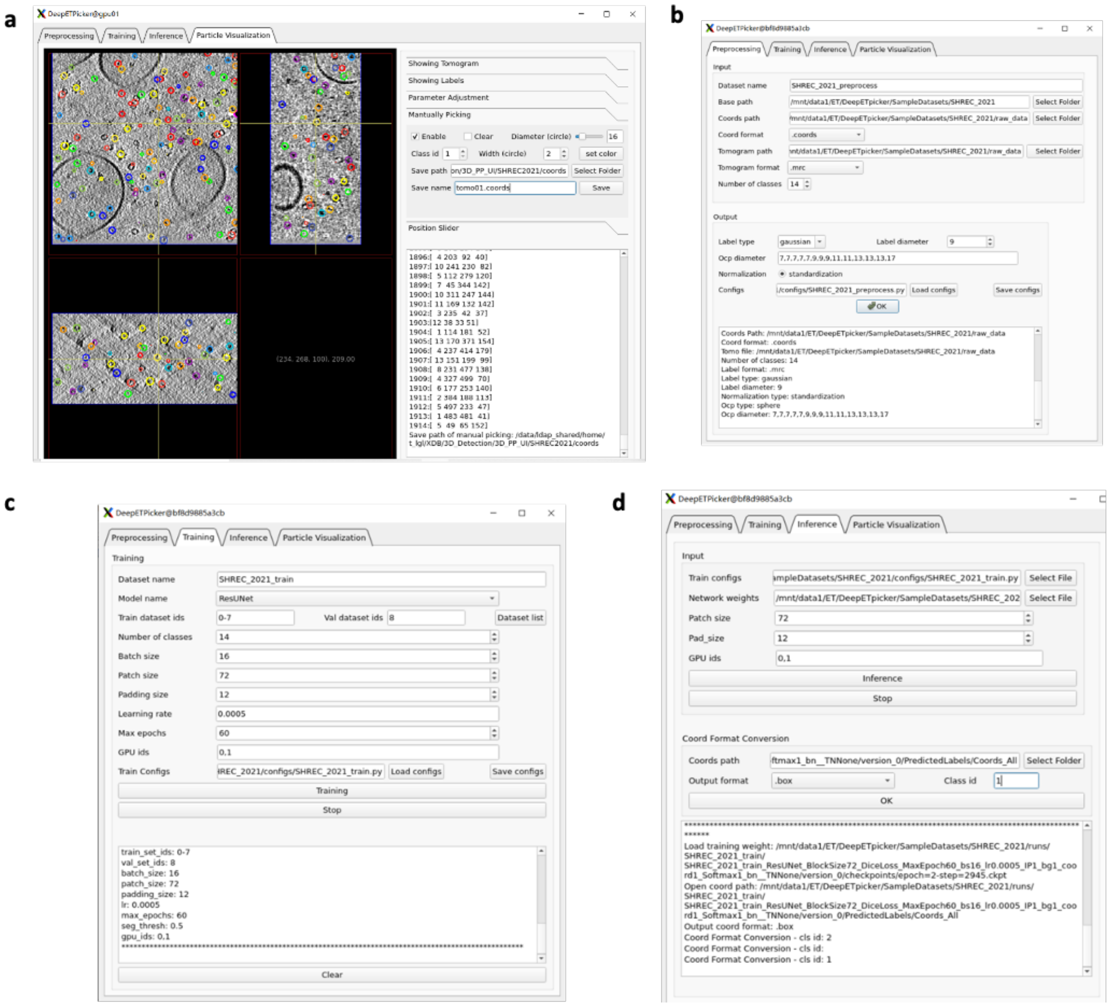


**Supplementary Fig. 1 | Graphical user interface of DeepETPicker. a**, The particle picking interface provides the following functions: manual annotation of particle centers and visualization of annotated particle centers. **b**, The pre-processing interface provides the following functions: pre-processing of raw cryo-ET tomograms and generation of simplified masks centered on annotated particle centers. **c**, The training interface provides the following functions: hyperparameter configuration and segmentation model selection. **d**, The inference interface provides the following functions: loading pretrained segmentation model, hyperparameter configuration, particle centers generation and format conversion for particle coordinates.

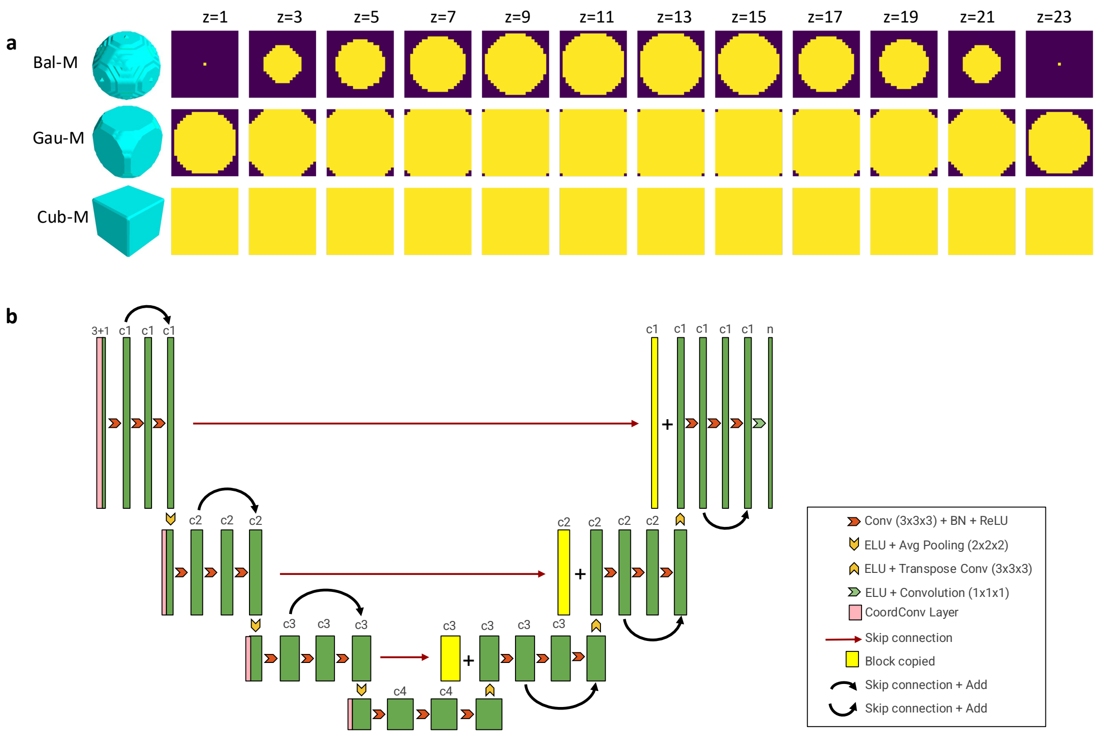


**Supplementary Fig. 2 | Illustration of simplified masks and network architecture of 3D-ResUNet. a,** Rendering of simplified/weak masks with a diameter $d = 23$ and their corresponding cross-sections at different positions $z$. Ball-M: ball masks. Gau-M: Gaussian masks. Cub-M: cubic masks. **b**, Detailed architecture of the segmentation neural network 3D-ResUNet, which is based on 3D-Unet[2] and adds residual connections of ResNet[3]. c1, c2, c3, c4 are feature map channels at different resolution levels.

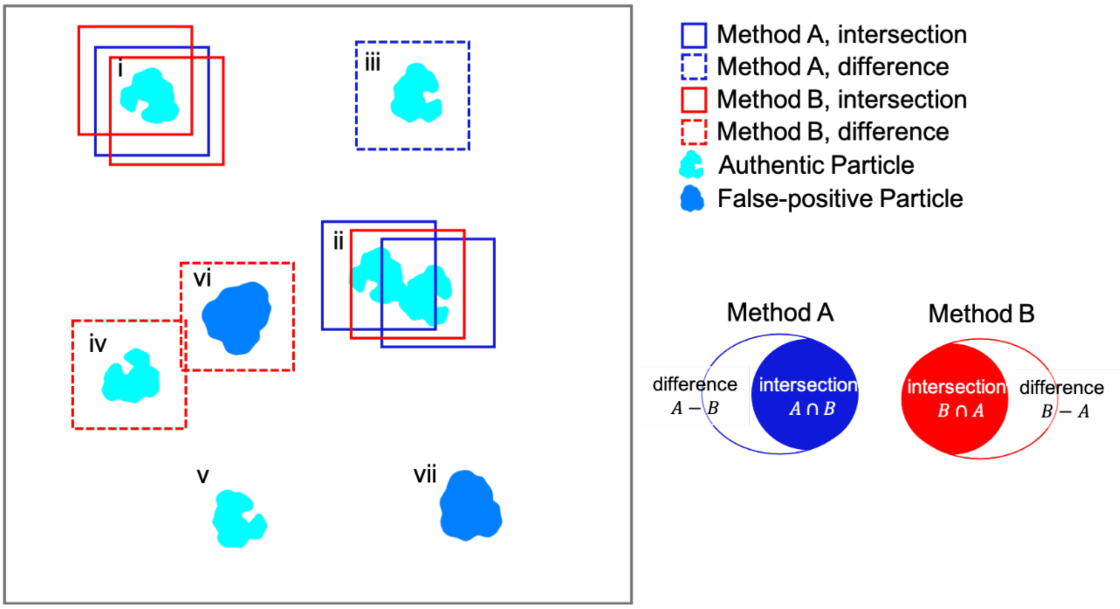


**Supplementary Fig. 3 | An explanation of intersection and difference sets of particles picked by methods A and B.** Solid boxes represent particles in the intersection set, and dashed boxes represent particles in the difference set. Particles picked by methods A and B are denoted in blue and red, respectively. Several representative cases of particles are shown. **Case i** shows a particle picked by both methods. Method A identifies it as a single particle, but method B identifies it as two separate particles with different centers. **Case ii** shows two particles in close proximity. Method A recognizes them as two separate particles with different centers, but method B recognizes it as a single particle. **Cases iii and iv** shows difference particles identified only by method A and method B, respectively. **Case v** shows particles missed by both methods. **Case vi** shows a false-positive particle similar in size or shape to the target particle identified by method B. **Case vii** shows a false-positive particle excluded by both methods. Either case i or ii may lead to different numbers of intersection sets between $A \cap B$ and $B \cap A$. Specifically, taking case i as an example, under the above definition of "same particles", the particle will count as 1 in A ∩ B and will count as 2 in B ∩ A. In summary, the seemingly counterintuitive asymmetry is caused the above definition of "same particles". Although this counter-intuitive asymmetry can be eliminated by modifying the definition of "same particles", the current way of setting $A \cap B$ and $B \cap A$ is maintained in this study because it carries useful information.

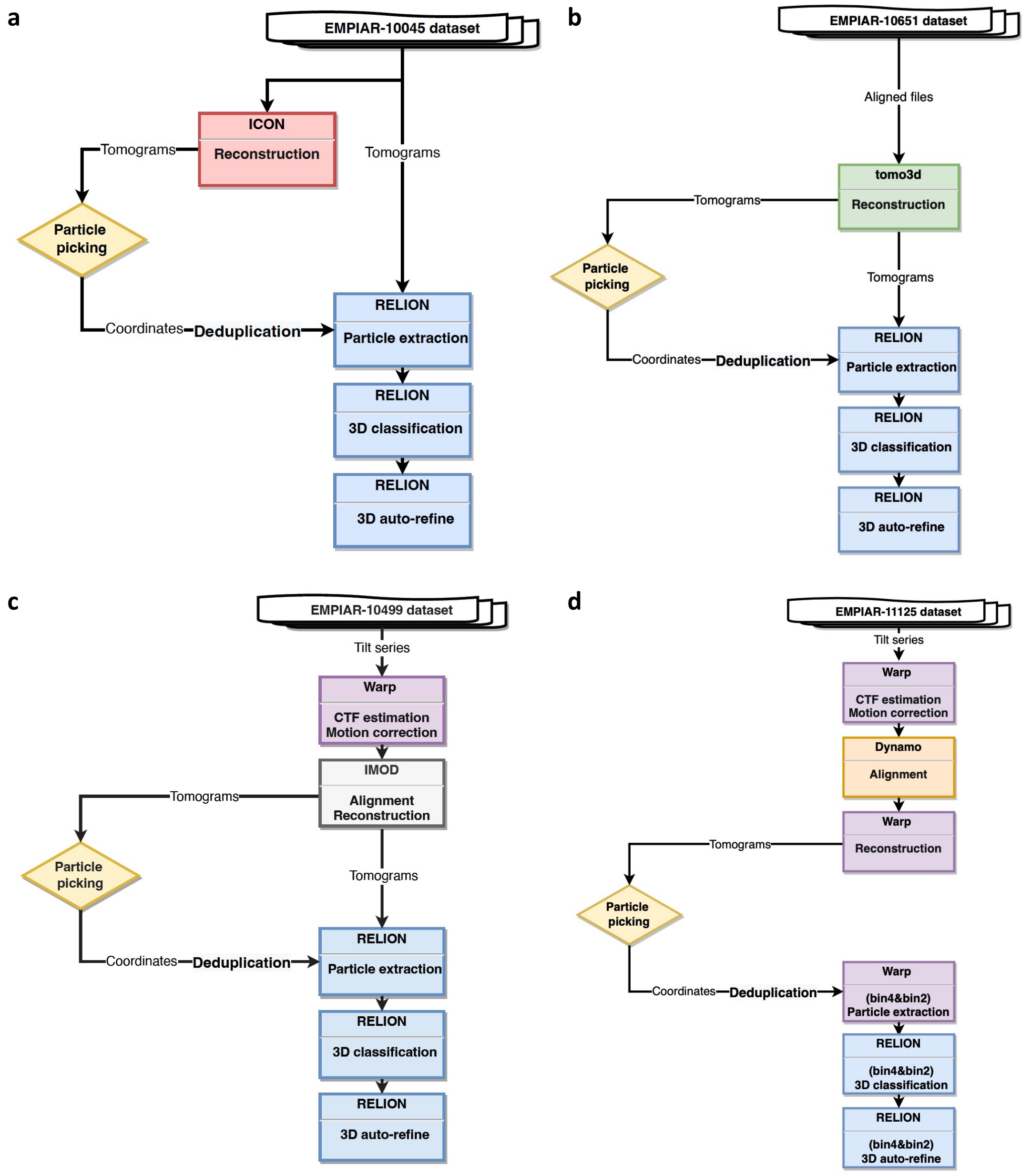


**Supplementary Fig. 4 | Sub-tomogram analysis workflow. a**, We use the aligned tilt series in the subdirectory of the EMPIAR entry to perform ICON reconstruction, which is then used for particle picking. The original tomograms of the entry are then utilized for sub-tomogram analysis, including CTF estimation, particle extraction, 2D classification, 3D auto-refine, and post-processing. The CTF model of each particle was generated by CTFFIND 4 in RELION 2.1.0. **b**, We use the aligned tilt series contained in subdirectory of the EMPIAR entry (EMPIAR-10651) to perform reconstruction by tomo3d (version: January 2015). Particle coordinates and the unbinned tomograms are utilized for sub-tomogram analysis by RELION 2.1.0, including CTF estimation, particle extraction, 3D classification, 3D auto-refinement. The CTF model of each particle was generated by CTFFIND 4 in RELION 2.1.0. **c**, Pre-processing operations of EMPIAR-10499 tilt series include motion correction and CTF estimation by Warp 1.0.9, tilt series alignment by IMOD 4.9.12, and reconstruction by weighted back projection in IMOD 4.9.12. After particle picking, all subsequent processes, including particle extraction, 3D classification, 3D auto-refine, and post-processing, are performed using RELION 2.1.0. The CTF model of each particle was generated by CTFFIND 4 in RELION 2.1.0. **d**, The pre-processing of EMPIAR-11125 tilt series include motion correction and CTF estimation by Warp 1.0.9, alignment by Dynamo v1.1.509_MCR-9.6.0, and reconstruction by Warp. After particle picking, the particle coordinates are used for particle extraction in Warp and the subsequent 3D classification and auto-refinement are performed using RELION 3.1 beta.

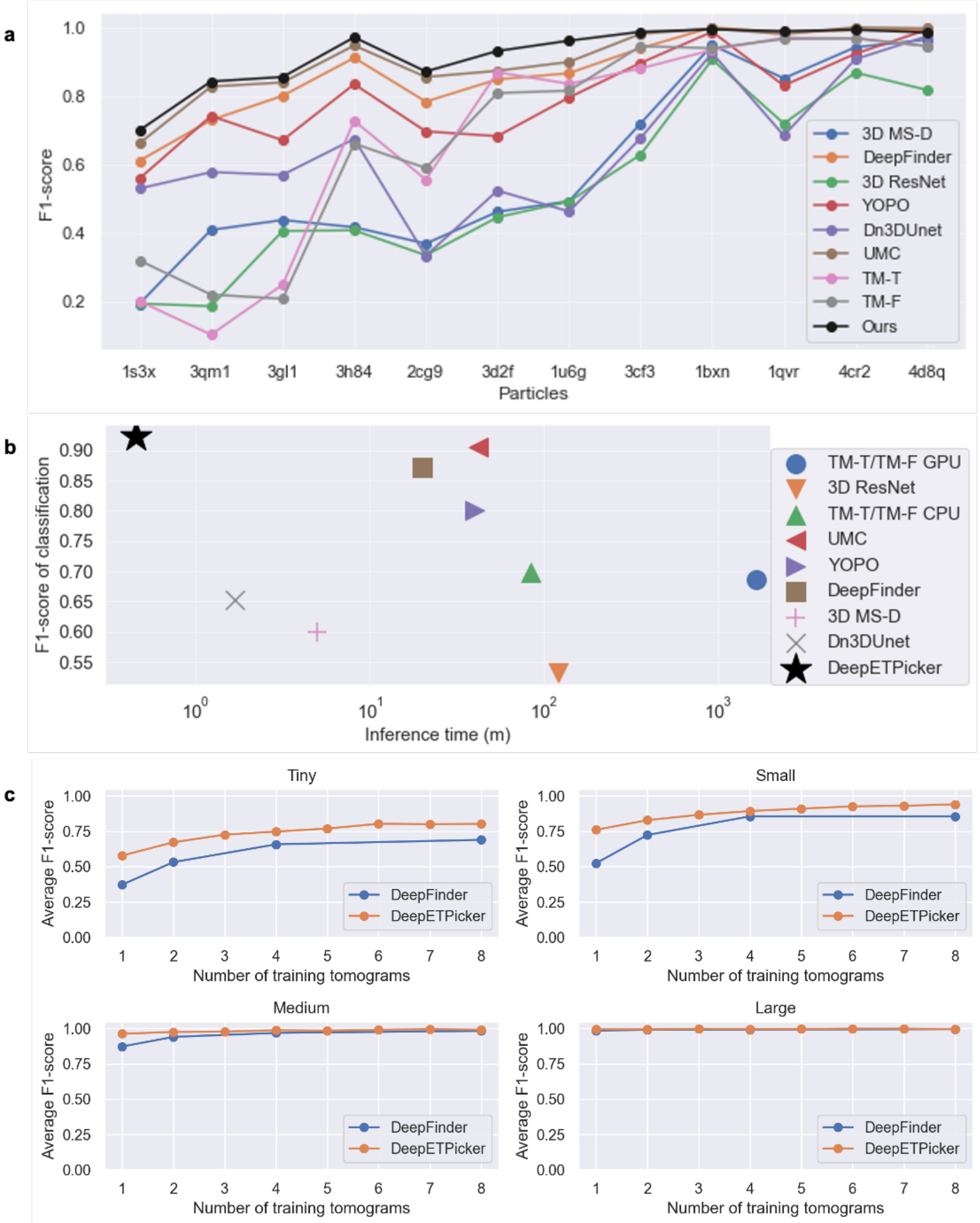


**Supplementary Fig. 5 | Particle picking performance of DeepETPicker in comparison with that of competing methods on the SHREC2020 dataset. a,** Classification performance measured in F1-score is plotted against particle molecular weight for DeepETPicker and other particle picking methods reported in the SHREC2020 challenge[4]. **b**, DeepETPicker runs substantially faster and achieves substantially higher classification performance than competing particle picking methods on the SHREC2020 dataset. Inference of DeepETPicker is performed on an Nvidia GeForce GTX 2080Ti. See Supplementary Table 6 for further details. **c**, Classification performance of DeepETPicker in comparison with DeepFinder measured in F1-

score plotted under different numbers of training tomograms and particles with different sizes on the SHREC2020 dataset. The particles are divided into four groups with different sizes, including Tiny (1s3x, 3qm1, 3gl1), Small (3h84, 2cg9, 3d2f, 1u6g), Medium (3cf3, 1bxn, 1qvr), and Large (4cr2, 4d8q).

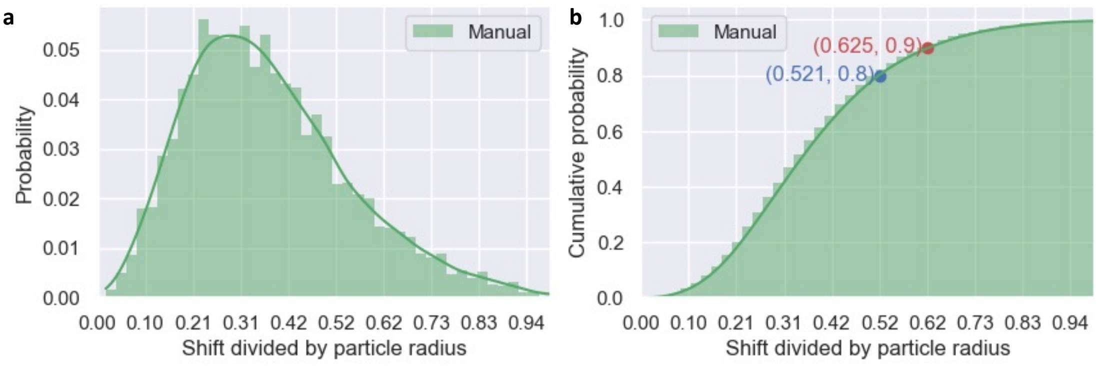


**Supplementary Fig. 6 | Euclidean distances between particle coordinates obtained by manual picking and particle coordinates after refinement, based on the dataset EMPIAR-10499:** Probability plot (a) and cumulative probability plot (b) of the distances, which are normalized by the particle radius.

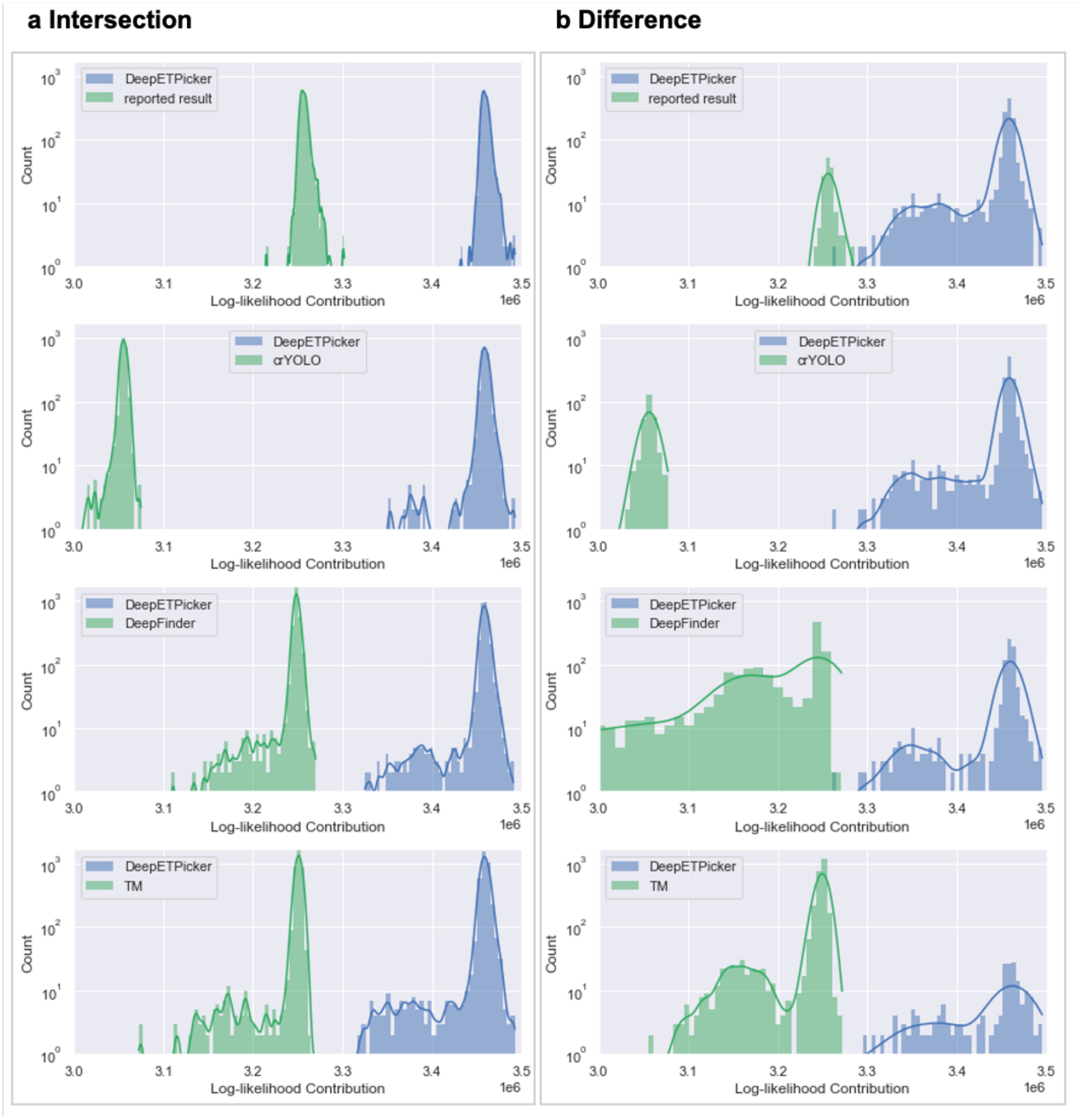


**Supplementary Fig. 7 | The log-likelihood contributions of the intersection (a) and difference (b) sets of particles between DeepETPicker and the other four methods on EMPIAR-10045 dataset.** The results of DeepETPicker and each of the other four methods are shown in blue and green, respectively.

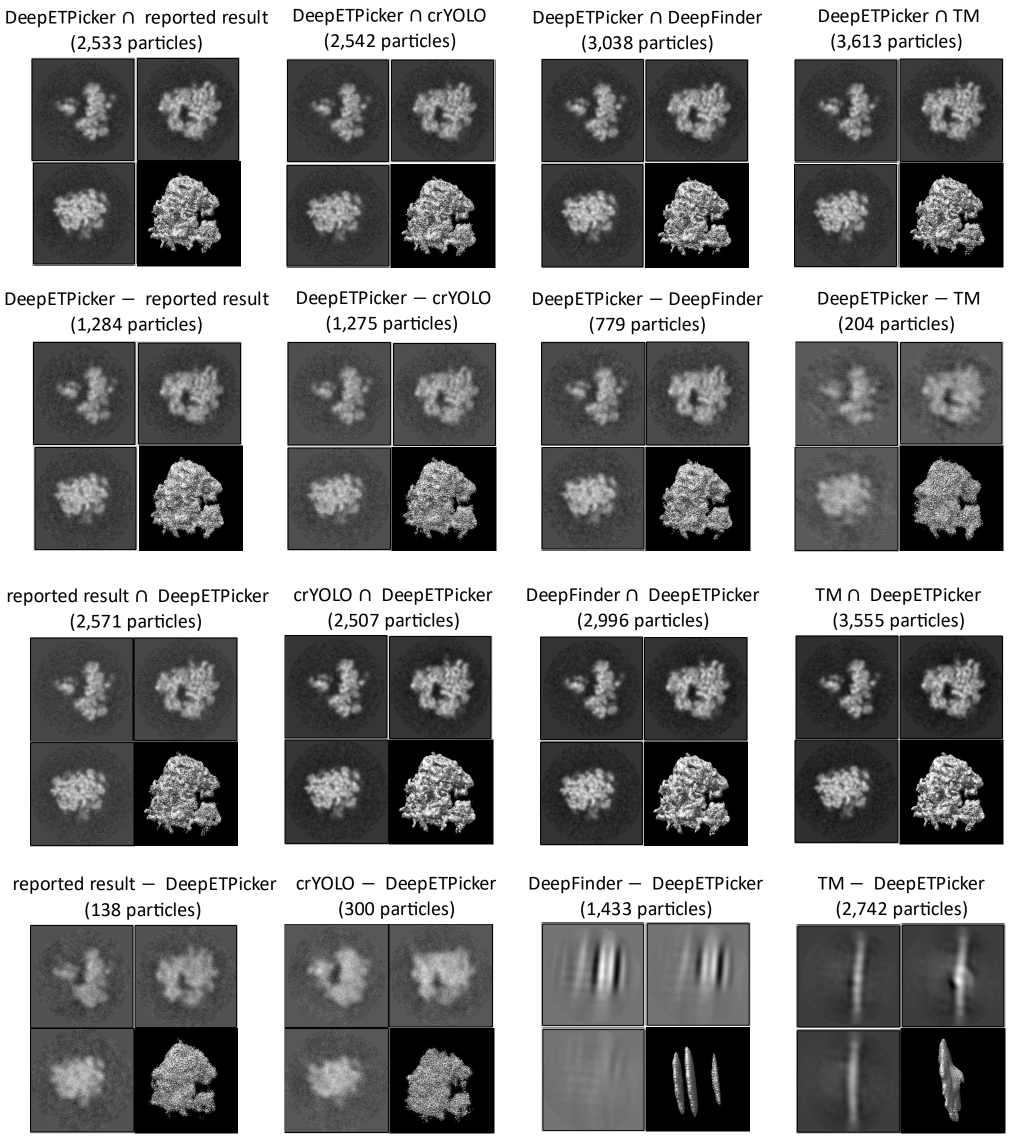


**Supplementary Fig. 8 | Comparison of sub-tomogram averaging for the intersection and difference sets of particles picked by different methods on the EMPIAR-10045 dataset of *S. cerevisiae* 80S ribosome**. The four figure panels are the 75$^{th}$, 95$^{th}$ and 125$^{th}$ sections of the sub-tomogram averaging result and the density map.

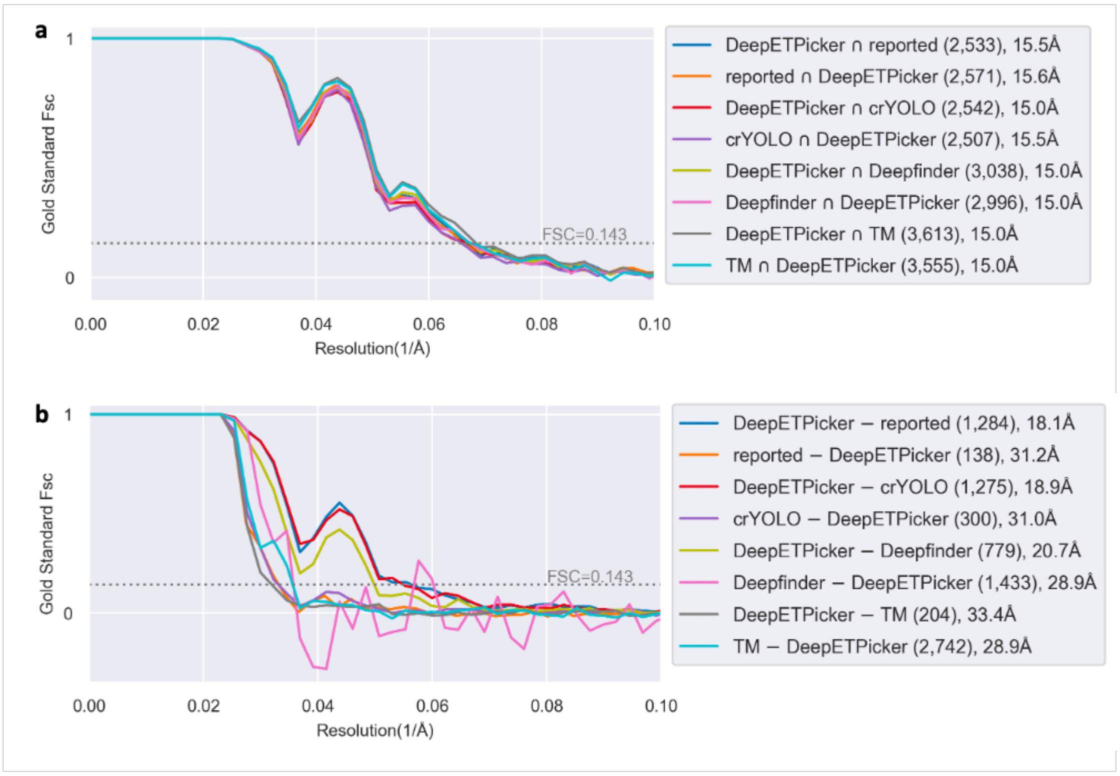


**Supplementary Fig. 9 | Comparison of FSC curves achieved by the intersection (a) and difference (b) sets of particles selected by different methods on the EMPIAR-10045 dataset of *S. cerevisiae* 80S ribosome.**

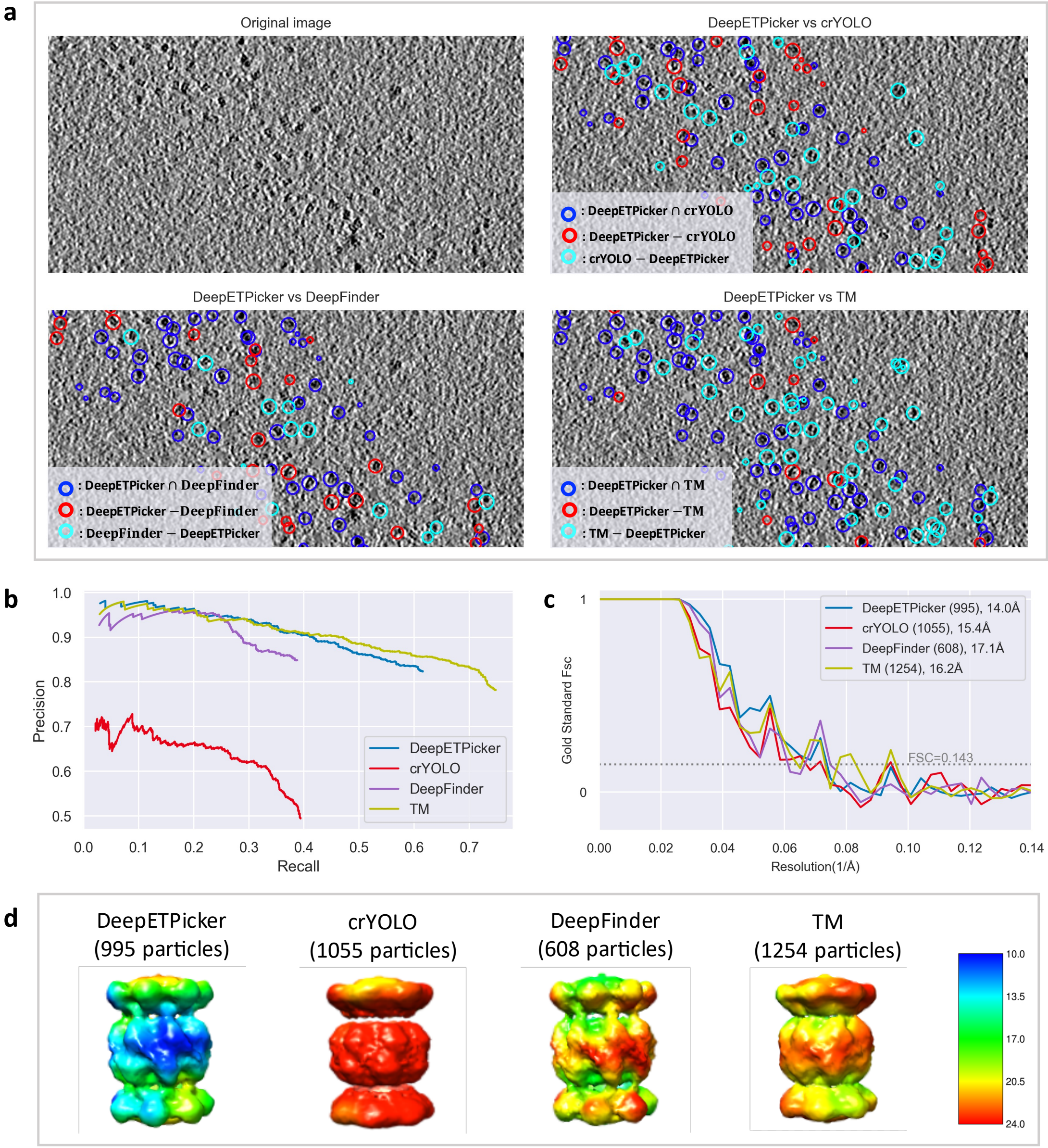


**Supplementary Fig. 10 | Particle picking results on the EMPIAR-10651 dataset**. **a**. Comparison of particle distributions between DeepETPicker and the other four methods (manual pick, crYOLO, template matching, and Deepfinder). Different color shows the intersection and differences of two particle sets. (This is a slice of $z = 100$ from the reconstruction of k2dft20s_14apra0023). Intersection set particles picked by DeepETPicker and the other method are shown in blue, difference set particles picked by DeepETPicker and the other method are shown in red and cyan, respectively. **b**. Precision-recall curves of different methods using manual particles as the reference. **c.** Comparative of gold standard FSC curves of particles picked by different methods on EMPIAR-10651 dataset. Of note, the oscillations of FSC curves are due to the small number of particles used.

**d.** Comparison of the local resolutions of sub-tomogram averages using different particle picking methods (DeepETPicker, crYOLO, Deepfinder and template matching).

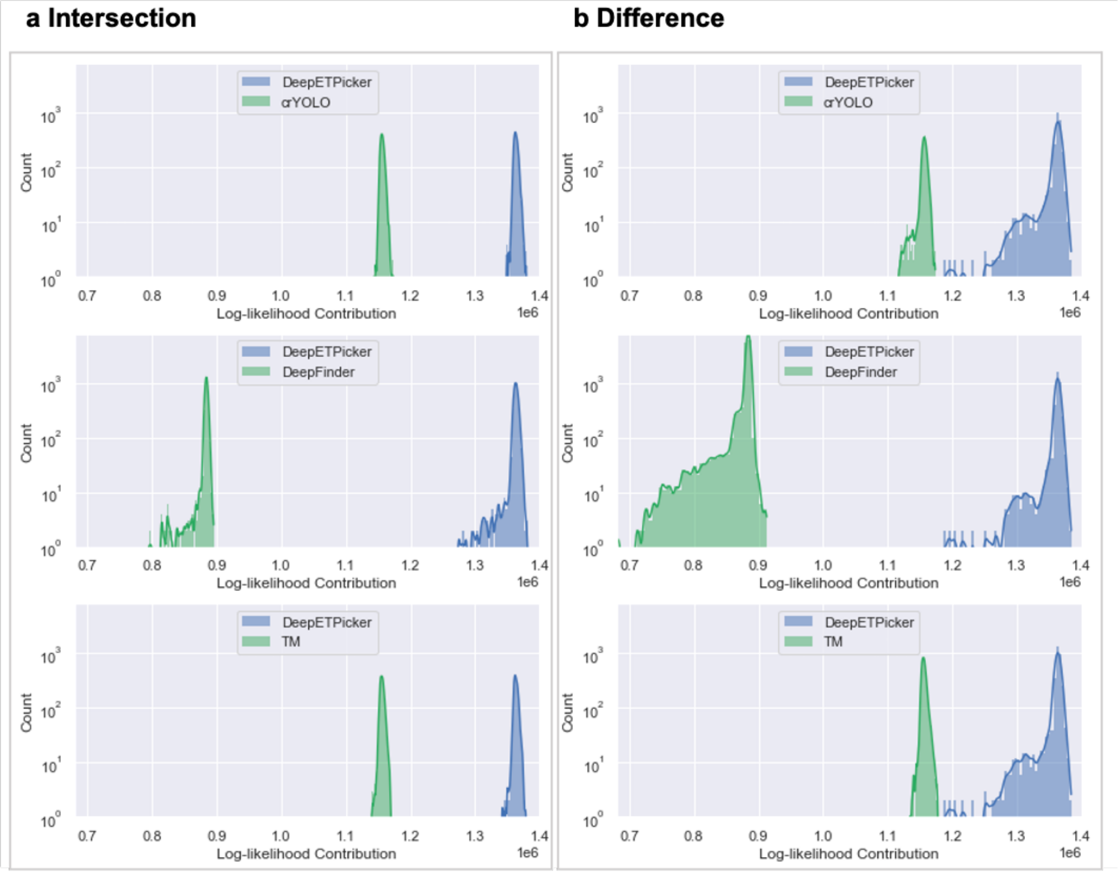


**Supplementary Fig. 11 | Log-likelihood contributions of the intersection (a) and difference (b) sets of particles between DeepETPicker and the other three methods on EMPIAR-10499 dataset.** The results of DeepETPicker and each of the other three competing methods are shown in blue and green, respectively.

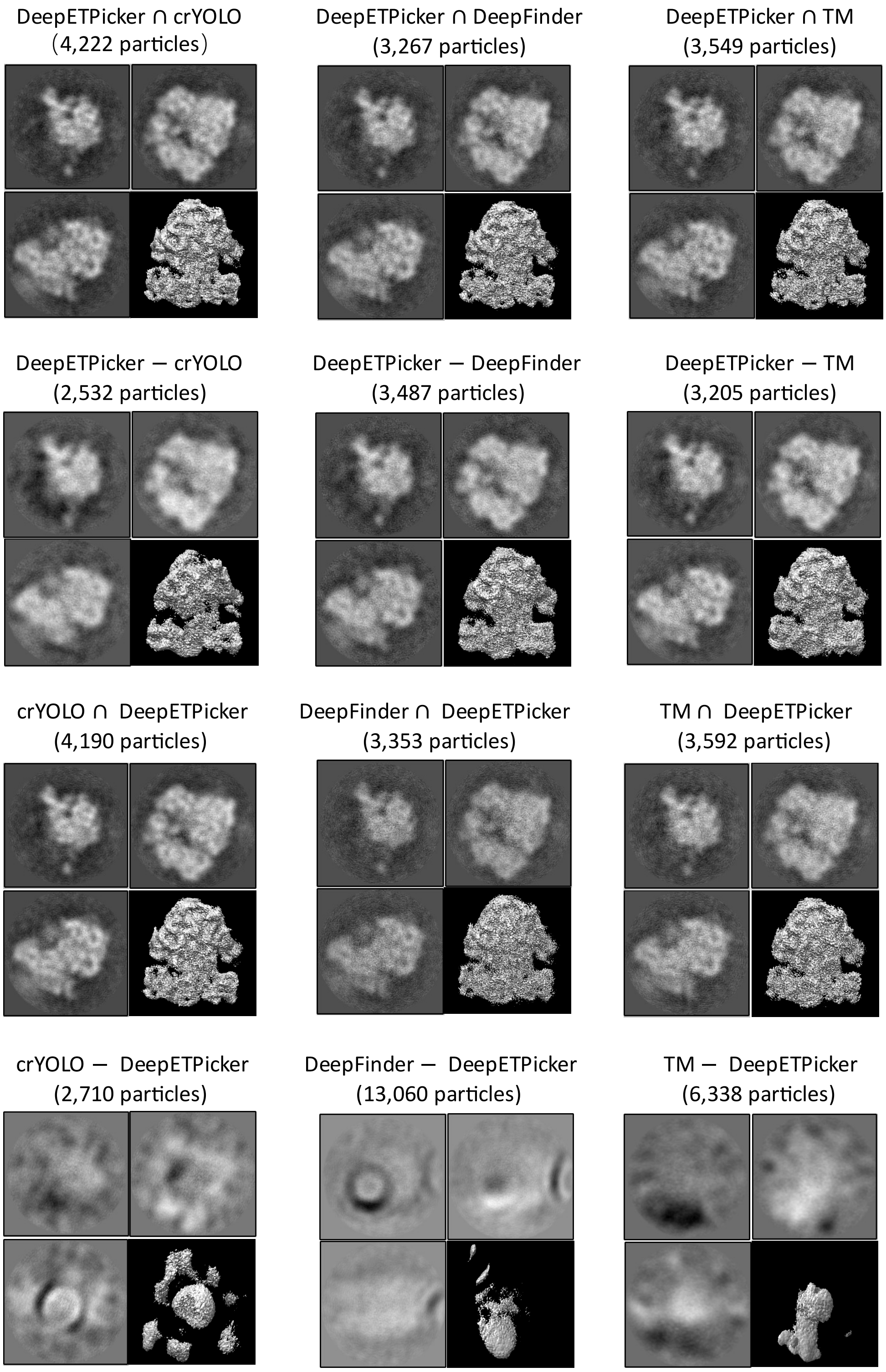


**Supplementary Fig. 12 | Comparison of subtomogram averaging for the intersection and difference sets of particles selected by different methods on EMPIAR-10499 dataset of native *M. pneumoniae* cells.** The four figure panels are the 75th, 95th and 125th sections of the sub-tomogram averaging result and the density map.

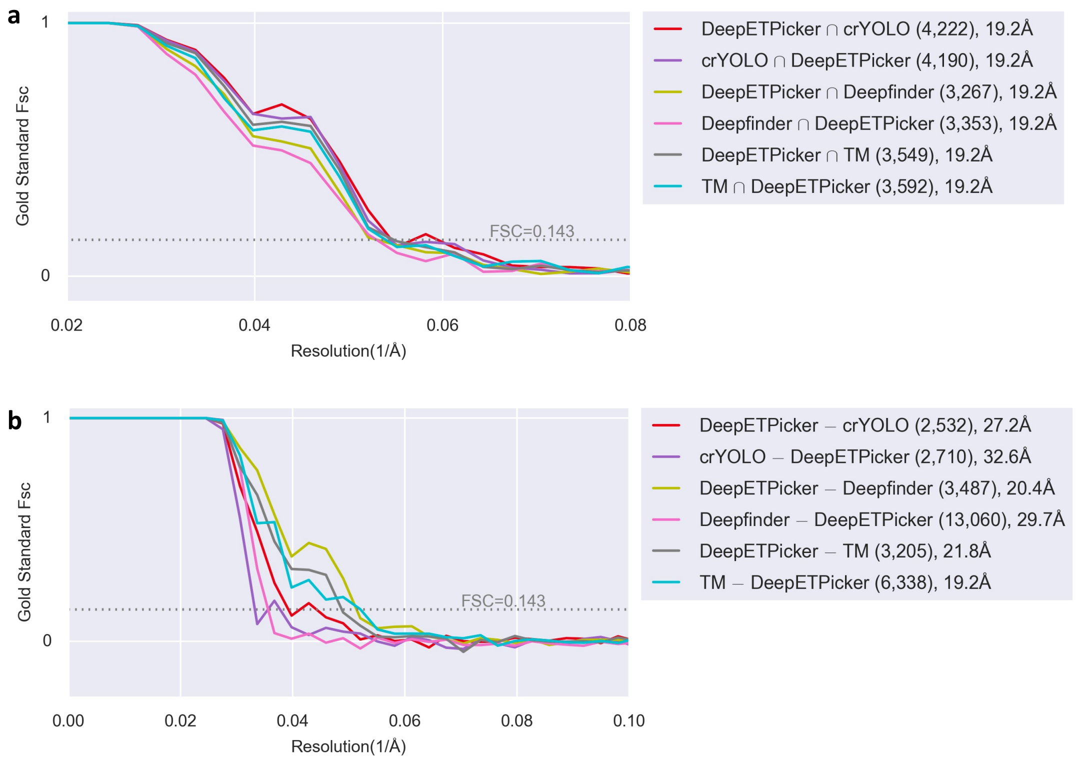


**Supplementary Fig. 13 | Comparison of FSC curves achieved by the intersection (a) and difference (b) sets of particles picked by different methods on EMPIAR-10499 dataset of native *M. pneumoniae* cells.**

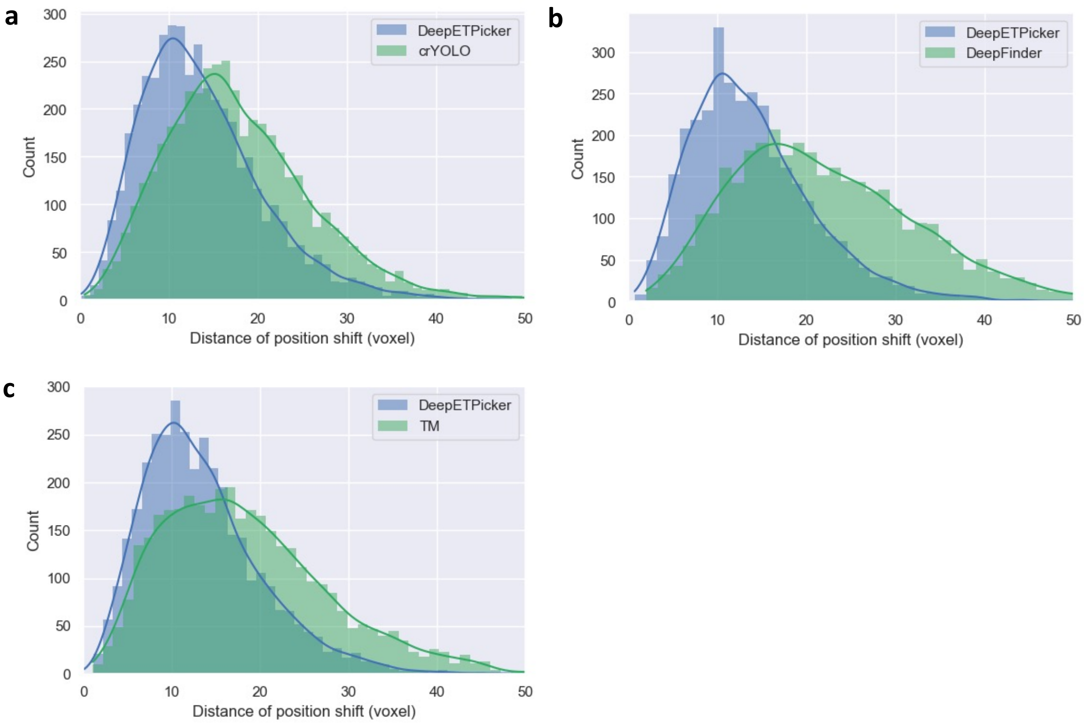


**Supplementary Fig. 14 | The two-norm of center shifts for the same particles picked by DeepETPicker and the competing methods on EMPIAR-10499 dataset.** (a) DeepETPicker vs crYOLO. (b) DeepETPicker vs DeepFinder. (c) DeepETPicker vs TM.

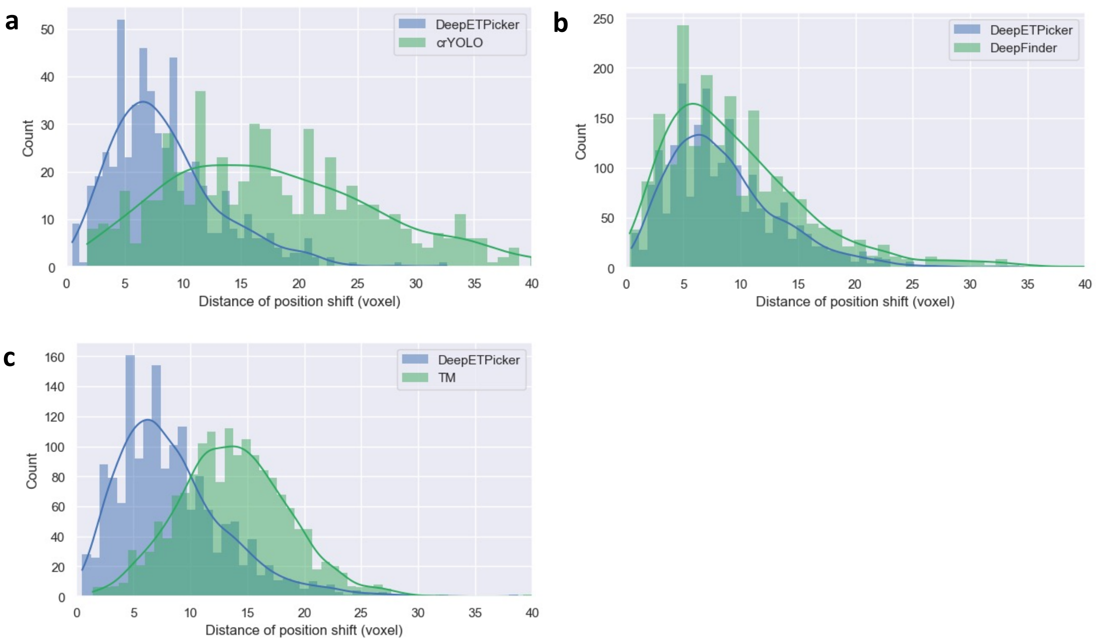


**Supplementary Fig. 15 | The two-norm of center shifts for the same particles picked by DeepETPicker and the competing methods on EMPIAR-11125 dataset.** (a) DeepETPicker vs crYOLO. (b) DeepETPicker vs DeepFinder. (c) DeepETPicker vs TM.

### C. Supplementary Tables

**Supplementary Table 1 | Hyperparameters of DeepETPicker on different datasets.**

| Datasets / Parameters | SHREC2020 | SHREC2021 | EMPIAR-10045 | EMPIAR-10499 | EMPIAR-10651 | EMPIAR-11125 | Parameter description |
|---|---|---|---|---|---|---|---|
| $t_g$ | 0.368 | | | | | | Threshold for Gaussian masks |
| $t_{seg}$ | 0.5 | | | | | | Threshold for transforming segmentation maps into binary maps |
| $d$ | 7~19 | 7~19 | 25 | 25 | 21 | 13 | Diameter of the particles |
| $t_{dist}$ | $\left\lceil \frac{d}{2} \right\rceil$ | | | | | | Threshold to distinguish whether two adjacent particles are the same particle, where $\lceil \cdot \rceil$ denotes the round-up operation |
| $t_{lm}$ | 0.1 | | | | | | Threshold for determine whether a local maximum is a particle |
| $N$ | 72 | 72 | 72 | 72 | 72 | 72 | Size of sub-tomograms |
| $pad\_size$ | 12 | 12 | 12 | 12 | 12 | 12 | Padding size for the overlap strategy |
| $lr$ | 0.001 | 0.001 | 0.001 | 0.001 | 0.001 | 0.001 | Initial learning rate |
| $batch\_size$ | 16 | 16 | 24 | 24 | 24 | 24 | Batch size |
| max_$epoch$ | 60 | 60 | 60 | 60 | 60 | 60 | Total number of training epochs |

**Supplementary Table 2 | Three types of simplified masks, i.e., Gaussian masks (Gau-M), cubic masks (Cub-M) and ball masks (Bal-M), with different diameters.** Size Based denotes the diameter of the generated mask is set as proportional to the size of real mask, Const7 and Const9 denote the diameters of the generated masks are set as 7 and 9, respectively.

| Simplified Mask | | 1s3x | 3qml | 3gl1 | 3h84 | 2cg9 | 3d2f | 1u6g | 3cf3 | 1bxn | 1qvr | 4cr2 | 5mrc | fiducial |
|---|---|---|---|---|---|---|---|---|---|---|---|---|---|---|
| **Gau-M** | Size Based | 7 | 7 | 7 | 7 | 9 | 9 | 9 | 11 | 13 | 13 | 13 | 17 | 7 |
| | Const7 | | | | | | | 7 | | | | | | |
| | Const9 | | | | | | | 9 | | | | | | |
| **Cub-M** | Size Based | 7 | 7 | 7 | 7 | 9 | 9 | 9 | 11 | 13 | 13 | 13 | 17 | 7 |
| | Const7 | | | | | | | 7 | | | | | | |
| | Const9 | | | | | | | 9 | | | | | | |
| **Bal-M** | Size Based | 7 | 7 | 7 | 7 | 9 | 9 | 9 | 11 | 13 | 13 | 13 | 17 | 7 |
| | Const7 | | | | | | | 7 | | | | | | |
| | Const9 | | | | | | | 9 | | | | | | |

**Supplementary Table 3 | Localization and classification performance of DeepETPicker trained by different types of simplified masks on the SHREC2021 dataset.**

| | | Localization | | | | | | | | | Classification | | | | | | | | | | | | | |
|---|---|---|---|---|---|---|---|---|---|---|---|---|---|---|---|---|---|---|---|---|---|---|---|---|
| **Mask Type** | | **RR** | **TP** | **FP** | **FN** | **MH** | **AD** | **R** | **P** | **F1** | **1s3x** | **3qm** | **3gl1** | **3h84** | **2cg9** | **3d2f** | **1u6g** | **3cf3** | **1bxn** | **1qvr** | **4cr2** | **5mrc** | **fiducial** | **mean F1** |
| **Real Mask** | | 1609 | 1479 | 70 | 86 | 59 | 1.47 | 0.94 | 0.92 | 0.93 | 0.39 | 0.52 | 0.58 | 0.81 | 0.80 | 0.83 | 0.79 | 0.96 | 0.99 | 0.95 | 0.97 | 0.99 | 1.00 | 0.81 |
| **Gau-M** | Size Based | 1510 | 1441 | 42 | 124 | 26 | 1.16 | 0.92 | 0.95 | 0.94 | 0.37 | 0.54 | 0.60 | 0.83 | 0.84 | 0.89 | 0.82 | 0.99 | 0.99 | 0.99 | 1.00 | 1.00 | 1.00 | 0.84 |
| | Const7 | 1510 | 1447 | 43 | 118 | 19 | 1.15 | 0.92 | 0.96 | 0.94 | 0.41 | 0.44 | 0.60 | 0.88 | 0.81 | 0.89 | 0.81 | 0.98 | 1.00 | 0.99 | 1.00 | 1.00 | 1.00 | 0.83 |
| | Const9 | 1508 | 1446 | 41 | 119 | 18 | 1.22 | 0.92 | 0.96 | 0.94 | 0.45 | 0.54 | 0.64 | 0.83 | 0.81 | 0.89 | 0.82 | 0.97 | 0.99 | 0.98 | 1.00 | 1.00 | 1.00 | 0.84 |
| **Cub-M** | Size Based | 1473 | 1421 | 33 | 144 | 19 | 1.22 | 0.91 | 0.97 | 0.93 | 0.33 | 0.53 | 0.60 | 0.85 | 0.76 | 0.87 | 0.77 | 0.97 | 0.98 | 0.97 | 1.00 | 1.00 | 1.00 | 0.82 |
| | Const7 | 1573 | 1466 | 82 | 99 | 24 | 1.13 | 0.93 | 0.93 | 0.93 | 0.46 | 0.55 | 0.59 | 0.83 | 0.83 | 0.88 | 0.83 | 0.99 | 1.00 | 0.96 | 1.00 | 1.00 | 1.00 | 0.84 |
| | Const9 | 1543 | 1438 | 71 | 127 | 31 | 1.22 | 0.92 | 0.93 | 0.92 | 0.35 | 0.47 | 0.48 | 0.82 | 0.77 | 0.85 | 0.82 | 1.00 | 1.00 | 0.98 | 1.00 | 1.00 | 1.00 | 0.81 |
| **Bal-M** | Size Based | 1497 | 1434 | 43 | 131 | 18 | 1.172 | 0.91 | 0.96 | 0.94 | 0.46 | 0.51 | 0.62 | 0.75 | 0.77 | 0.81 | 0.81 | 0.96 | 0.99 | 0.97 | 1.00 | 1.00 | 1.00 | 0.82 |
| | Const7 | 1284 | 1202 | 73 | 363 | 9 | 1.192 | 0.77 | 0.94 | 0.84 | 0.47 | 0.46 | 0.61 | 0.83 | 0.81 | 0.83 | 0.80 | 0.97 | 0.99 | 0.97 | 0.00 | 0.00 | 1.00 | 0.67 |
| | Const9 | 1474 | 1418 | 43 | 147 | 11 | 1.214 | 0.90 | 0.96 | 0.93 | 0.36 | 0.53 | 0.63 | 0.88 | 0.81 | 0.87 | 0.80 | 0.99 | 1.00 | 0.98 | 1.00 | 1.00 | 1.00 | 0.84 |

**Supplementary Table 4 | Comparison of DeepETPicker versus competing methods in localization performance**. The methods such as URFinder, DeepFinder, U-CLSTM, MC-DS-Net, YOPO, TM-F and TM are reported in the SHREC2021 challenge[5]. RR: detected number of particles; TP: true positive; FP: false positive, FN: false negative, MH: multiple hits, AD: average Euclidean distance from predicted particle center in voxels; Recall: uniquely selected true locations divided by actual number of particles in the test tomogram; Precision: uniquely selected true locations divided by RR; Miss rate: percentage of results that yield negative results; F1 Score: harmonic average of the precision and recall. The best results in each column are highlighted in bold. ↑ indicates that the higher the better, ↓ indicates that the lower the better.

| Methods | RR | TP↑ | FP↓ | FN↓ | MH↓ | AD↓ | Recall↑ | Precision↑ | Miss rate↓ | F1 Score↑ |
|---|---|---|---|---|---|---|---|---|---|---|
| URFinder | 1969 | 1298 | 377 | 267 | 149 | 1.84 | 0.826 | 0.659 | 0.174 | 0.733 |
| YOPO | 1627 | 1224 | 232 | 341 | **14** | 1.66 | 0.720 | 0.752 | 0.221 | 0.765 |
| CFN | 1765 | 1364 | 239 | 201 | 20 | 1.52 | 0.868 | 0.773 | 0.132 | 0.818 |
| U-CLSTM | 1460 | 1253 | 49 | 312 | 44 | 2.13 | 0.798 | 0.858 | 0.202 | 0.827 |
| MC DS Net | 1760 | 1415 | 239 | 150 | 56 | 1.59 | 0.901 | 0.804 | 0.099 | 0.850 |
| DeepFinder | 1567 | 1362 | 64 | 203 | 20 | 2.22 | 0.867 | 0.869 | 0.133 | 0.868 |
| DeepETPicker | 1510 | **1447** | **43** | **118** | 19 | **1.15** | **0.921** | **0.958** | **0.079** | **0.939** |

**Supplementary Table 5 | Comparison of DeepETPicker versus competing methods in classification performance measured in F1-score on specific particle classes.** The competing methods such as URFinder, DeepFinder, U-CLSTM, MC-DS-Net, YOPO, TM-F and TM are reported in SHREC2021 challenge[5]. The best results in each column are highlighted in bold.

| Methods | 1s3x | 3qm1 | 3gl1 | 3h84 | 2cg9 | 3d2f | 1u6g | 3cf3 | 1bxn | 1qvr | 4cr2 | 5mrc | fiducial | mean F1 |
|---|---|---|---|---|---|---|---|---|---|---|---|---|---|---|
| URFinder | **0.00** | **0.42** | **0.45** | **0.60** | **0.54** | **0.67** | **0.67** | **0.87** | **0.97** | **0.86** | **0.93** | **0.95** | **0.43** | **0.64** |
| YOPO | 0.20 | 0.15 | 0.47 | 0.60 | 0.63 | 0.63 | 0.61 | 0.88 | 0.94 | 0.92 | 0.98 | 0.97 | 0.95 | 0.69 |
| U-CLSTM | 0.28 | 0.42 | 0.39 | 0.56 | 0.51 | 0.65 | 0.57 | 0.95 | 0.99 | 0.90 | 0.99 | **1.00** | **1.00** | 0.71 |
| DeepFinder | 0.40 | 0.48 | 0.52 | 0.70 | 0.72 | 0.77 | 0.74 | 0.96 | 0.99 | 0.95 | 0.97 | 1.00 | **1.00** | 0.78 |
| CFN | 0.25 | **0.51** | **0.61** | 0.77 | 0.71 | 0.76 | 0.73 | 0.97 | **1.00** | 0.97 | 1.00 | **1.00** | **1.00** | 0.79 |
| MC DS Net | 0.32 | 0.49 | 0.60 | 0.78 | 0.78 | 0.79 | 0.80 | 0.96 | 0.99 | 0.93 | 0.98 | **1.00** | **1.00** | 0.80 |
| DeepETPicker | **0.41** | 0.44 | 0.60 | **0.88** | **0.81** | **0.89** | **0.81** | **0.98** | **1.00** | **0.99** | **1.00** | **1.00** | **1.00** | **0.83** |

**Supplementary Table 6 | Reported computing time in training and inference stages for processing one SHREC2021 tomogram of size $200 \times 512 \times 512$.** Calculation of estimated speedup ratios of DeepETPicker in the inference stage is based on the assumption that the calculation is mainly completed by the GPU.

| Methods | Training stage | Inference stage | Hardware | FP32 TFLOPS | Estimated speedup ratio of DeepETPicker in inference stage |
|---|---|---|---|---|---|
| URFinder | 300h | 2h6m | Nvidia Quodro RTX 8000 GPU (2×) | 16.3 x 2 | 654.42 |
| DeepFinder | 50h | 20m | Nvidia M40 (1×) | 6.83 | 21.76 |
| U-CLSTM | 120h | 15m | Nvidia Quodro RTX-5000 GPU (1×) | 11.2 | 26.77 |
| MC DS Net | 22h | 5m | Nvidia GeForce RTX 3090 GPU (1×) | 35.58 | 28.34 |
| YOPO | 8h | 40m | Nvidia GeForce Titan X GPU (1×) | 6.69 | 42.63 |
| CFN | 96h | - | Nvidia GeForce RTX 3090 GPU (2×) | 35.58 x 2 | - |
| TM-F/TM GPU | N/A | 4h26m | Nvidia GeForce GTX 1080Ti (1×) | 11.34 | 480.58 |
| DeepETPicker | 17h | 28s | Nvidia GeForce GTX 2080Ti (1×) | 13.45 | 1.00 |

**Supplementary Table 7 | Relationship between standard deviation of Gaussian kernels and SNR levels of the SHREC2021 dataset.**

| Standard deviation of Gaussian kernel | SNR level of SHREC2021 datasets |
|---|---|
| 0 | 0.127-0.587 |
| 0.5 | 0.116-0.535 |
| 1.1 | 0.101-0.463 |
| 1.5 | 0.077-0.348 |
| 2 | 0.056-0.254 |
| 3 | 0.039-0.171 |
| 5 | 0.026-0.110 |

**Supplementary Table 8 | Ablation Study of DeepETPicker**. RC=Residual Connection, CC=Coord Conv, IP=Image Pyramid, DA=Data Augmentation, DD=De-Duplication, OT=Overlap-tile Strategy.

| Ablation Study | | | | | | Localization | | | | | | | | | Classification | | | | | | | | | | | | | |
|---|---|---|---|---|---|---|---|---|---|---|---|---|---|---|---|---|---|---|---|---|---|---|---|---|---|---|---|---|
| RC | CC | IP | DA | DD | OT | RR | TP | FP | FN | MH | AD | R | P | F1 | 1s3x | 3qm1 | 3gl1 | 3h84 | 2cg9 | 3d2f | 1u6g | 3cf3 | 1bxn | 1qvr | 4cr2 | 5mrc | fiducial | mean F1 |
| | | | | | | 1353 | 1221 | 17 | 344 | 106 | 1.40 | 0.78 | 0.90 | 0.84 | 0.20 | 0.36 | 0.37 | 0.57 | 0.59 | 0.67 | 0.58 | 0.85 | 0.98 | 0.87 | 0.98 | 0.97 | 1.00 | 0.69 |
| ✓ | | | | | | 1295 | 1197 | 21 | 368 | 71 | 1.38 | 0.76 | 0.92 | 0.84 | 0.21 | 0.33 | 0.39 | 0.62 | 0.61 | 0.67 | 0.61 | 0.86 | 0.97 | 0.89 | 0.97 | 0.97 | 1.00 | 0.70 |
| ✓ | ✓ | ✓ | | | | 1528 | 1286 | 28 | 279 | 184 | 1.49 | 0.82 | 0.84 | 0.83 | 0.36 | 0.37 | 0.39 | 0.59 | 0.61 | 0.68 | 0.66 | 0.85 | 0.97 | 0.85 | 0.97 | 0.95 | 1.00 | 0.71 |
| ✓ | ✓ | ✓ | ✓ | | | 1424 | 1314 | 35 | 251 | 72 | 1.27 | 0.84 | 0.92 | 0.88 | 0.38 | 0.42 | 0.56 | 0.77 | 0.72 | 0.83 | 0.77 | 0.93 | 1.00 | 0.92 | 0.98 | 0.99 | 1.00 | 0.79 |
| ✓ | ✓ | ✓ | ✓ | ✓ | | 1376 | 1313 | 33 | 252 | 28 | 1.27 | 0.84 | 0.95 | 0.89 | 0.37 | 0.42 | 0.54 | 0.77 | 0.73 | 0.83 | 0.77 | 0.95 | 1.00 | 0.93 | 0.98 | 0.99 | 1.00 | 0.79 |
| ✓ | ✓ | ✓ | ✓ | ✓ | ✓ | 1510 | 1447 | 43 | 118 | 19 | 1.15 | 0.92 | 0.96 | 0.94 | 0.41 | 0.44 | 0.60 | 0.88 | 0.81 | 0.89 | 0.81 | 0.98 | 1.00 | 0.99 | 1.00 | 1.00 | 1.00 | 0.83 |

**Supplementary Table 9 | Comparison of DeepETPicker versus competing methods in global resolution achieved by the intersection and difference sets of particles selected by different methods on EMPIAR-10045 dataset of S. cerevisiae 80S ribosome.** RH resolution is the theoretical resolution estimated based on the Rosenthal and Henderson B-factor plot (RH plot)[1]

| Method | No of particles for 3D reconstruction | Resolution (Å) | RH Resolution (Å) | Method | No of particles for 3D reconstruction | Resolution (Å) | RH Resolution (Å) |
|---|---|---|---|---|---|---|---|
| DeepETPicker ∩ reported result | 2533 | 15.5 | 15.7 | reported result ∩ DeepETPicker | 2571 | 15.6 | - |
| DeepETPicker − reported result | 1284 | 18.1 | 17.8 | reported result − DeepETPicker | 138 | 31.2 | - |
| DeepETPicker ∩ crYOLO | 2542 | 15.0 | 15.7 | crYOLO ∩ DeepETPicker | 2507 | 15.5 | 16.0 |
| DeepETPicker − crYOLO | 1275 | 18.9 | 17.8 | crYOLO − DeepETPicker | 300 | 31.0 | 24.7 |
| DeepETPicker ∩ DeepFinder | 3038 | 15.0 | 15.3 | DeepFinder ∩ DeepETPicker | 2996 | 15.0 | 16.4 |
| DeepETPicker − DeepFinder | 779 | 20.7 | 19.9 | DeepFinder − DeepETPicker | 1433 | 28.9 | 18.8 |
| DeepETPicker ∩ TM | 3613 | 15.0 | 14.9 | TM ∩ DeepETPicker | 3555 | 15.0 | 16.2 |
| DeepETPicker − TM | 204 | 33.4 | 35.6 | TM − DeepETPicker | 2742 | 28.9 | 17.0 |

**Supplementary Table 10 | Comparison of DeepETPicker versus competing methods in global resolution achieved by the intersection and difference sets of particles selected by different methods on EMPIAR-10499 dataset of native M. pneumoniae cells.** RH resolution is the theoretical resolution estimated based on the Rosenthal and Henderson B-factor plot (RH plot)[1]

| Method | No of particles picked & selected for 3D reconstruction | Resolution (Å) | RH Resolution (Å) | Method | No of particles picked & selected for 3D reconstruction | Resolution (Å) | RH Resolution (Å) |
|---|---|---|---|---|---|---|---|
| DeepETPicker ∩ crYOLO | 4222 | 19.2 | 19.3 | crYOLO ∩ DeepETPicker | 4190 | 19.2 | 20.3 |
| DeepETPicker − crYOLO | 2532 | 27.2 | 20.6 | crYOLO − DeepETPicker | 2710 | 32.6 | 21.4 |
| DeepETPicker ∩ DeepFinder | 3267 | 19.2 | 20.0 | DeepFinder ∩ DeepETPicker | 3353 | 19.2 | 34.0 |
| DeepETPicker − DeepFinder | 3487 | 20.4 | 19.8 | DeepFinder − DeepETPicker | 13060 | 29.7 | 29.5 |
| DeepETPicker ∩ TM | 3549 | 19.2 | 19.8 | TM ∩ DeepETPicker | 3592 | 19.2 | 22.9 |
| DeepETPicker − TM | 3205 | 21.8 | 20.0 | TM − DeepETPicker | 6338 | 19.2 | 20.9 |

### References


1 Rosenthal, P. B. & Henderson, R. Optimal determination of particle orientation, absolute hand, and contrast loss in single-particle electron cryomicroscopy. *Journal of Molecular Biology* **333**, 721-745 (2003).

2 Çiçek, Ö., Abdulkadir, A., Lienkamp, S. S., Brox, T. & Ronneberger, O. 3D U-Net: learning dense volumetric segmentation from sparse annotation. in *International Conference on Medical Image Computing and Computer-Assisted Intervention (MICCAI).* 424-432, Springer, (2016).

3 He, K., Zhang, X., Ren, S. & Sun, J. Deep residual learning for image recognition. in *IEEE Conference on Computer Vision and Pattern Recognition (CVPR).* 770-778, (2016).

4 Gubins, I. *et al.* SHREC 2020: classification in cryo-electron tomograms. *Computers & Graphics* **91**, 279-289 (2020).

5 Gubins, I. *et al.* SHREC 2021: classification in cryo-electron tomograms. in *Eurographics Workshop on 3D Object Retrieval.* (eds Silvia Biasotti *et al.*), The Eurographics Association, (2021).

### D. Supplementary Videos

**SupplementaryVideo01.avi** Comparison of particles picked by DeepETPicker versus the other four competing methods (reported result: the method reported in the original article (Bharat & Scheres, Nature Protocols, 11:2054-2065, 2016), crYOLO, template matching, and Deepfinder) on the EMPIAR-10045 experimental dataset. Different colors show the same and different particles detected. Intersection sets of particles picked by DeepETPicker and the other competing method are shown blue. Difference sets of particles picked by DeepETPicker and the other competing method are shown in red and cyan, respectively.

**SupplementaryVideo02.avi** Comparison of particles picked by DeepETPicker versus the other three competing methods (crYOLO, Deepfinder, and template matching) on the EMPIAR-10499 experimental dataset. Different colors show the same and different particles detected. Intersection sets of particles picked by DeepETPicker and the other competing method are shown in blue. Difference sets of particles picked by DeepETPicker and the other competing method are shown in red and cyan, respectively.
